# Computational Analysis Revealed Five Novel Mutations in Human *IL2RG* gene Related to X-SCID

**DOI:** 10.1101/528349

**Authors:** Tamadur Babiker Abbas Mohammed, Asma Ali Hassan Ali, Areeg ElsirAbdelgadir Elemam, Wala Omer Mohammed Altayb, Tebyan Ameer Abdelhameed Abbas, Mohamed Ahmed Salih Hassan

## Abstract

**Background:** X linked severe combined immunodeficiency (X-SCID) is a life-threatening disorder. It is due to mutation of the interleukin two receptor gamma-chain (IL2RG) gene. Nonsynonymous SNPs (nsSNPs) are the most common polymorphism, known to be deleterious or disease-causing variations because they alter protein sequence, structure, and function. Objective: is to reveal the effect of harmful SNPs in the function and structure of IL2RG protein.

**Method:** Data on IL2RG was investigated from dbSNP/NCBI database. Prediction of damaging effect was done using sift, polyphen, provean and SNAP2.more software were used for more analysis: phd-snp, and and go, Pmut, Imutant.modeling was done using chimera and project hope. Gene interaction was done by gene mania.3UTR prediction was done using polymiRTS software.

**Result:** The in-silico prediction identified 1479 SNPs within IL2RG gene out of which 253 were coding SNPs, 50 took place in the miRNA 3 UTR, 21 occurred in 5 UTR region and 921 occurred in intronic regions. a total of 12 missense nsSNPs were found to be damaging by both a sequence homology-based tool (SIFT) and a structural homology-based method (PolyPhen), Five of them were novel; rs1322436793(<u>G305R</u>), rs1064794027(<u>C182Y</u>), rs111033620(<u>G114D</u>), rs193922347 (<u>Y105C</u>) and rs1293196743(<u>Y91C</u>), Two SNPs(<u>Rs144075871</u> and <u>rs191726889</u>) out of 50 in the 3UTR region were predicted to disrupt miRNAs binding sites and affect the gene expression.

**Conclusions:** Computational analysis of SNPs has become a very valuable tool in order to discriminate neutral SNPs from damaging SNPs. This study revealed 5 novel nsSNPs in the IL2RG gene by using different software and 21 SNPs in 3UTR. These SNPs could be considered as important candidates in causing diseases related to IL2RG mutation and could be used as diagnostic markers.

## 1. Introduction

Severe combined immunodeficiency (SCID) [OMIM: #300400] is a rare life-threatening condition characterized by impaired cellular and humoral immunity ^(1)^. X linked severe combined immunodeficiency (X-SCID; 308380) is the commonest type of SCID (∼50%) ^(2-5)^. Interleukin 2 receptor gamma-chain gene (*IL2RG)* mutation is responsible of X-SCID ^(6-9)^. Interleukin 2 receptor gamma-chain gene (*IL2RG)* [OMIM: *308380] is positioned on chromosome Xq13 and its product is common gamma protein ^(10-12)^. The common gamma is an elemental structure of the receptors for IL-2, IL-4, IL-7, IL-9, IL-15, and IL-21^(13, 14)^.These signaling cytokines are essential for lymphocytes development and function, they promotes regulation of T-cell growth, differentiation and peripheral tolerance, increasing of natural killer (NK) cytolytic activity and B cells differentiation^(6, 8, 15)^. T cell dysfunction effects immunoglobulin class switching of B cells that mean patients have compromised specific antibody responses, so they have defects in class-switched immunoglobulins (IgG, IgA, and IgE) with detectable IgM ^(3)^. In classical X-SCID nonfunctional *IL2RG* results in deficiency of T and natural killer (NK) lymphocytes and malfunctioning B lymphocytes T (-), NK (-) B (+). non-classical X-SCID (312863) may have a missense mutation or other potentially non-loss of function result is low numbers of T cells while NK and B cells are normal in numbers T (low) NK (+) B (+) ^(16-18)^. Male patients with classical X-SCID suffer from severe opportunistic infections and they usually die during infancy if not treated while non-classical X-SCID has less severe type of disease ^(19-21)^

There are large numbers of information about *IL2RG* mutations associated with X-SCID found in the literatures ^(4, 16, 22, 23)^. Moreover, there are many publications covering polymorphism of *IL2RG* single nucleotide polymorphism (SNPs) like: D39N, G114D, C115F, C115R, I153N, L162H, C182R, L183S, R222C, R224W, R226H, W240C, S241I, R285Q, L293Q ^(12, 16, 17, 23-27)^. Also, a large numbers of missenses mutations or nonsynonymous (SNP) of *IL2RG* are collected in international databases. Despite this there are insufficient studies investigating all the harmful nsSNPs of *IL2RG*. The aim of this study to the possible effect of *IL2RG* SNPs on function and structure. Single Nucleotide Polymorphisms (SNPs) are variation of DNA sequence in which a single nucleotide is alter. It is the most common polymorphism ^(28)^. It can occur in coding or non-coding DNA sequence, SNPs is detected in the coding region of people leading to genetic variation ^(28,29)^. Non-synonymous SNPs (ns SNPs) are present in coding region of genome which is frequently leads to alteration in amino acid residues of gene product. This single lost or additional nucleotide causes a frame shift mutation ^(30)^. This effect changes the protein that is expressed, this possibly will cause a harmful mutation ^(31)^. This is the first study investigate SNPs located in *IL2RG* gene using computational approach.

## 2. Materials and methods

Data on *IL2RG* gene was obtained from national center for biological information (NCBI) web site (https://www.ncbi.nlm.nih.gov/) and the SNPs (single nucleotide polymorphisms) information was retrieved from NCBI SNPs database dbSNP (https://www.ncbi.nlm.nih.gov/snp/) (32). The gene ID and sequence was obtained from Uiprot (https://www.uniprot.org/). Analysis of the SNPs was done according to (figure 1).

**Figure 1:**
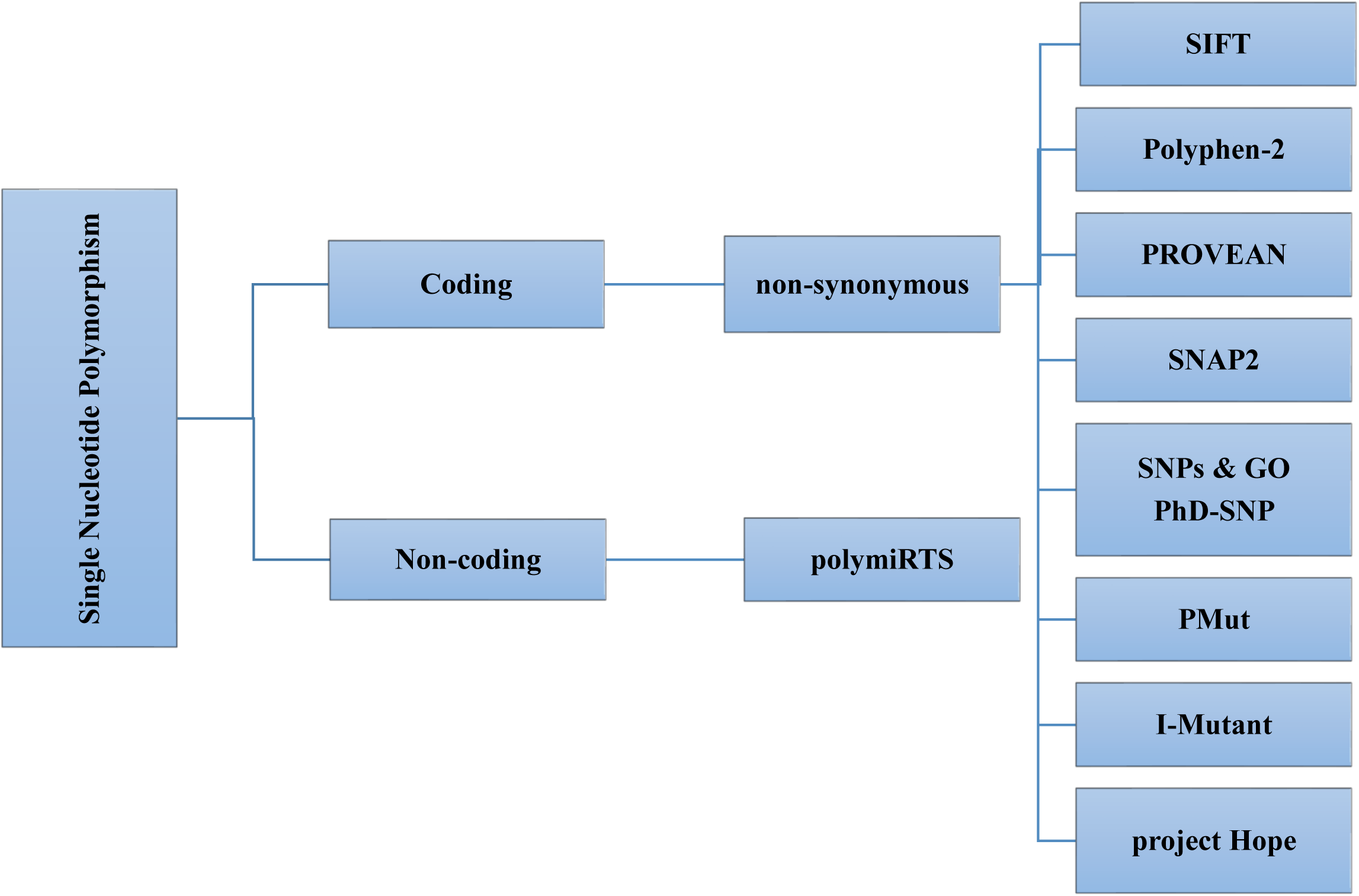
Flowchart of the software using in the analysis processes.

### 2.1. SIFT: Sorting Intolerant from Tolerant

software is available at (http://sift.bii.a-star.edu.sg/). SIFT is a sequence homology-based tool that sorts intolerant from tolerant amino acid substitutions and predicts whether an amino acid substitution in a protein will have a phenotypic outcome, considering the position at which the mutation occurred and the type of amino acid change. At the point when a protein sequence submitted, SIFT picks related proteins and acquired an alignment of these proteins with the query. According to the change on the type of amino acids appearing at each position in the alignment, SIFT calculates the probability that the substitution is tolerated or not. If this standardized esteem is less than a cutoff, the substitution is predicted to be deleterious. SIFT scores <0.05 are predicted to be intolerant or deleterious amino acid substitutions, whereas scores >0.05 are considered tolerant. ^(33)^

### 2.2. PolyPhen-2

(http://genetics.bwh.harvard.edu/pph2/) **Poly**morphism **Phen**otyping **v2.** It is a tool that predicts conceviable impact of an amino acid ulteration on the structure and function of a human protein by using simple physical and comparative considerations. The submission of the sequence allows querying for a single individual amino acid substitution or a coding, non-synonymous SNP commented on in the SNP database. This software calculates position-specific independent count (PSIC) scores for each of the two variants and calculate the difference of the PSIC scores of the two variants. The higher a PSIC score difference, the higher functional effect to have. PolyPhen scores were designated as probably damaging (0.95–1), possibly damaging (0.7–0.95), and benign (0.00–0.31). ^(34)^ ^(35)^

### 2.3. Provean

**Pr**otein **Va**riation **E**ffect **An**alyzer software available at (http://provean.jcvi.org/index.php) it predict whether an amino acid substitution has an impact on the biological function of protein. Provean is useful in filtrating sequence variants to distinguish non-synonymous variants that are predicted to be practically imperative. ^(36)^

### 2.4. SNAP2

is a trained classifier that is based on a machine learning gadget called “neural network”. It distinguishes between effect and neutral variants/non-synonymous SNPs by taking differences of sequence and variant features in considerations. Most important input for the prediction is the evolutionary information taken from multiple sequence alignment. Also, structural features such as predicted secondary structure and solvent accessibility are considered. If available also annotation of the sequence or close homologs are used in. SNAP2 has persistent two-state accuracy (effect/neutral) of 82%. Software is available at (https://rostlab.org/owiki/index.php/Snap2).^(37, 38)^

### 2.5. PHD-SNP

prediction of human Deleterious Single Nucleotide Polymorphisms is accessible at (http://snps.biofold.org/phd-snp/phd-snp.html). It is a Support Vector Machines (SVMs) based method trained to predict disease-associated nsSNPs using sequence information. The related transformaion is predicted as disease-related (Disease) or as neutral polymorphism (Neutral). ^(39)^

### 2.6. SNP & GO

is a server for the prediction of single point protein mutations likely to be involved in the causing of diseases in humans. SNP&GO is accessible at (https://snps-and-go.biocomp.unibo.it/snps-and-go/). ^(40)^

### 2.7. PMUT

available at (http://mmb.irbbarcelona.org/PMut/) is based on the use of different types of sequence information to detect mutations, and neural networks to analyse this information. It provides a very simple output: yes/no answer and a reliability index. ^(41)^

### 2.8. I-MUTANT

available at (http://gpcr.biocomp.unibo.it/∼emidio/I-Mutant3.0/old/IntroI-Mutant3.0_help.html) is a suit of support vector machine based predictors which integrated in a unique web server. It offers the chance to predict protein stability changes upon single site mutations starting from protein sequence alone or protein structure if accessible. Also, it gives opportunity to predict human deleterious SNPs from the protein sequence alone. ^(42)^

### 2.9. Project HOPE

online software. It is a web service where the user can submit a sequence and mutation. The software gathers basic data from different sources, including calculations on the 3D protein structure, sequence commented in UniProt and prediction from other software. It gathers this information to give analysis for the effect of a certain mutation on the protein structure. HOPE will show the effect of that mutation in such a way that any one even those without a bioinformatics background can understand it. It allows the user to submit a protein sequence or an accession number of the protein of interest. In a next step the user can choose the mutated amino acid with a simple mouse click. In the final step the user can simply tap on one of the other 19 amino acid types that will become the mutant residue, and then full report well be available. HOPE is available at (http://www.cmbi.ru.nl/hope/method/). ^(43)^

### 2.10. UCSF Chimera

(university of California at san Francisco) (https://www.cgl.ucsf.edu/chimera/) is an exceedingly extensible program for interactive visualization and analysis of molecular structures and relative data, including density maps, supramolecular assemblies, sequence alignments, docking results, directions, and conformational troupes. High-quality images and animations can be generated. **Chimera** includes complete documentation and several tutorials. **Chimera** is developed by the <u>Resource for Biocomputing, Visualization, andInformatics</u> (RBVI), supported in part by the <u>National Institutes of Health</u> (P41-GM103311).^(44)^

### 2.11. PolymiRTS

used to predict 3UTR (un-translated region): polymorphism in microRNAs and their target sites available at (http://compbio.uthsc.edu/miRSNP/) it is a data base of naturally occurring DNA variations in microRNAs(miRNA) seed region and miRNA target sites. MicroRNAs combine to the transcript of protein coding genes and cause translational repression or mRNA destabilization. SNPs in microRNA and their target sites may influence miRNA-mRNA interaction, causing impact on miRNA-mediated gene repression. PolymiRTS database was made by examining 3UTRs of mRNAs in human and mouse for SNPs in miRNA target destinations. Then, the impact of polymorphism on gene expression and phenotypes are identified and then connected in the database. The PolymiRTS data base also includes polymorphism in target sites that have been supported by variety of experimental methods and polymorphism in miRNA seed regions. ^(45, 46)^

### 2.12. Gene interaction **GeneMAINIA**

software that finds other genes that are related to a set of input genes, using a very large set of functional association data. Association data include protein and genetic interactions, pathways, co-expression, co-localization and protein domain similarity. GeneMANIA also used to find new members of a pathway or complex, find additional genes you may have missed in your screen or find new genes with a specific function, such as protein kinases. This software is available at (https://genemania.org/).^(47)^

## 3. Results

### 3.1. Retrieval of SNPs from the Database

A total of 1479 SNPs within *IL2RG* gene -at the time of the study-were retrieved from dbSNP database, out of which 253 were coding SNPs, 50 took place in the miRNA 3′ UTR, 21 occurred in 5′ UTR region and 921 occurred in intronic regions.

**Figure (2).**
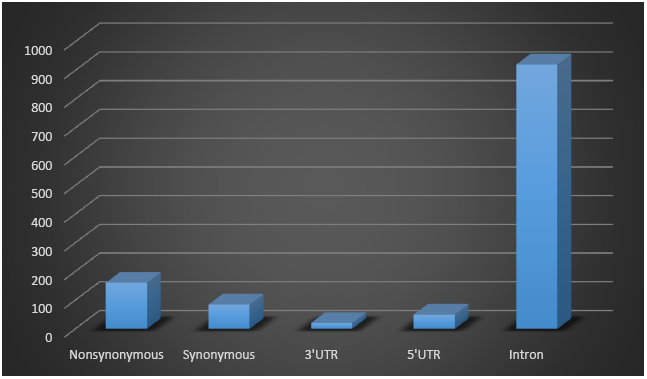
distribution of nonsynonymous, synonymous, 3′ UTR, 5′UTR and intronic SNPs for *IL2RG* retrieved from NCBI database.

### 3.2.

**Table (1).** Prediction of functional effect of Deleterious and damaging nsSNPs by SIFT, Polyphen-2, PROVEAN and SNAP2:

| **Chr.**<br>**Position\*** | **dbSNP ID** | **Amino**<br>**Acid**<br>**Change** | **SIFT Prediction** | **SIFT**<br>**score** | **Polyphen-2**<br>**Prediction** | **Polyphen-2**<br>**Score** | **PROVEAN**<br>**Prediction** | **PROVEAN**<br>**score** | **SNAP2**<br>**Prediction** | **SNAP2**<br>**Score** |
| --- | --- | --- | --- | --- | --- | --- | --- | --- | --- | --- |
| **71108288** | rs1322436793 | G305R | Deleterious | 0 | probably damaging | 1 | Deleterious | -3.457 | effect | 76 |
| **71109267** | rs1057517950 | W240R | Deleterious | 0 | probably damaging | 1 | Deleterious | -13.448 | effect | 91 |
| **71109309** | rs869320659 | R226C | Deleterious | 0.01 | probably damaging | 1 | Deleterious | -6.126 | effect | 49 |
| **71109315** | rs869320658 | R224W | Deleterious | 0 | probably damaging | 1 | Deleterious | -7.601 | effect | 90 |
| **71109321** | rs111033618 | R222C | Deleterious | 0.03 | probably damaging | 1 | Deleterious | -4.007 | effect | 72 |
| **71110205** | rs1064794027 | C182Y | Deleterious | 0.02 | probably damaging | 1 | Deleterious | -10.504 | effect | 55 |
| **71110506** | rs137852511 | L151P | Deleterious | 0.02 | probably damaging | 1 | Deleterious | -3.865 | effect | 19 |
| **71110615** | rs111033622 | C115R | Deleterious | 0 | probably damaging | 1 | Deleterious | -11.55 | effect | 99 |
| **71110617** | rs111033620 | G114D | Deleterious | 0.04 | probably damaging | 1 | Deleterious | -6.538 | effect | 98 |
| **71110644** | rs193922347 | Y105C | Deleterious | 0.05 | probably damaging | 1 | Deleterious | -8.563 | effect | 43 |
| **71110686** | rs1293196743 | Y91C | Deleterious | 0.01 | probably damaging | 1 | Deleterious | -7.039 | effect | 47 |
| **71110906** | rs1057520293 | L87P | Deleterious | 0.01 | probably damaging | 1 | Deleterious | -5.396 | effect | 83 |

\*Chr: chromosome

### 3.3.

**Table (2).** prediction of Disease Related and pathological effect of nsSNPs by PhD-SNP, SNPs & GO and PMut:

**3.3. Table (2) prediction of Disease Related and pathological effect of nsSNPs by PhD-SNP, SNPs & GO and PMut:**
| **Chr.**<br>**Position\*** | **dbSNP ID** | **Amino**<br>**Acid**<br>**Change** | **SNP & GO**<br>**Prediction** | **RI\*** | **Probability** | **PhD-SNP**<br>**Prediction** | **RI\*** | **Probability** | **PMUT Prediction** |  |
| --- | --- | --- | --- | --- | --- | --- | --- | --- | --- | --- |
| **71108288** | rs1322436793 | G305R | Disease | 5 | 0.727 | Disease | 6 | 0.793 | **0.76 (88%)** | Disease |
| **71109267** | rs1057517950 | W240R | Disease | 6 | 0.795 | Disease | 3 | 0.643 | **0.81 (89%)** | Disease |
| **71109309** | rs869320659 | R226C | Disease | 1 | 0.528 | Disease | 4 | 0.7 | **0.79 (89%)** | Disease |
| **71109315** | rs869320658 | R224W | Disease | 6 | 0.807 | Disease | 7 | 0.857 | **0.87 (92%)** | Disease |
| **71109321** | rs111033618 | R222C | Disease | 0 | 0.509 | Disease | 4 | 0.709 | **0.66 (85%)** | Disease |
| **71110205** | rs1064794027 | C182Y | Disease | 6 | 0.823 | Disease | 8 | 0.917 | **0.73 (87%)** | Disease |
| **71110506** | rs137852511 | L151P | Disease | 1 | 0.546 | Disease | 4 | 0.707 | **0.61 (83%)** | Disease |
| **71110615** | rs111033622 | C115R | Disease | 6 | 0.779 | Disease | 6 | 0.791 | **0.90 (93%)** | Disease |
| **71110617** | rs111033620 | G114D | Disease | 3 | 0.661 | Disease | 5 | 0.758 | **0.75 (87%)** | Disease |
| **71110644** | rs193922347 | Y105C | Disease | 3 | 0.627 | Disease | 4 | 0.697 | **0.79 (89%)** | Disease |
| **71110686** | rs1293196743 | Y91C | Disease | 2 | 0.599 | Disease | 3 | 0.672 | **0.76 (88%)** | Disease |
| **71110906** | rs1057520293 | L87P | Disease | 2 | 0.586 | Disease | 5 | 0.726 | **0.87 (91%)** | Disease |

\* Chr: chromosome \* RI: reliability index

### 3.4.

**Table (3).** Prediction of nsSNPs Impact on Protein structure Stability by I-Mutant:

**3.4. Table (3) Prediction of nsSNPs Impact on Protein structure Stability by I-Mutant:**
| **Chr.**<br>**Position\*** | **dbSNP ID\*** | **Amino Acid**<br>**Change** | **IMUTANT**<br>**PREDICTION** | **SCORE** |
| --- | --- | --- | --- | --- |
| **71108288** | rs1322436793 | G305R | DECREASE | -0.56 |
| **71109267** | rs1057517950 | W240R | DECREASE | -0.71 |
| **71109309** | rs869320659 | R226C | DECREASE | -1.27 |
| **71109315** | rs869320658 | R224W | DECREASE | -0.64 |
| **71109321** | rs111033618 | R222C | DECREASE | -1.4 |
| **71110205** | rs1064794027 | C182Y | DECREASE | -0.48 |
| **71110506** | rs137852511 | L151P | DECREASE | -1.49 |
| **71110615** | rs111033622 | C115R | INCREASE | 0.03 |
| **71110617** | rs111033620 | G114D | DECREASE | -0.94 |
| **71110644** | rs193922347 | Y105C | DECREASE | -1.05 |
| **71110686** | rs1293196743 | Y91C | DECREASE | -1.23 |
| **71110906** | rs1057520293 | L87P | DECREASE | -1.65 |

\* Chr: chromosome. \*db: database.

### 3.5.

**Figure 3:**
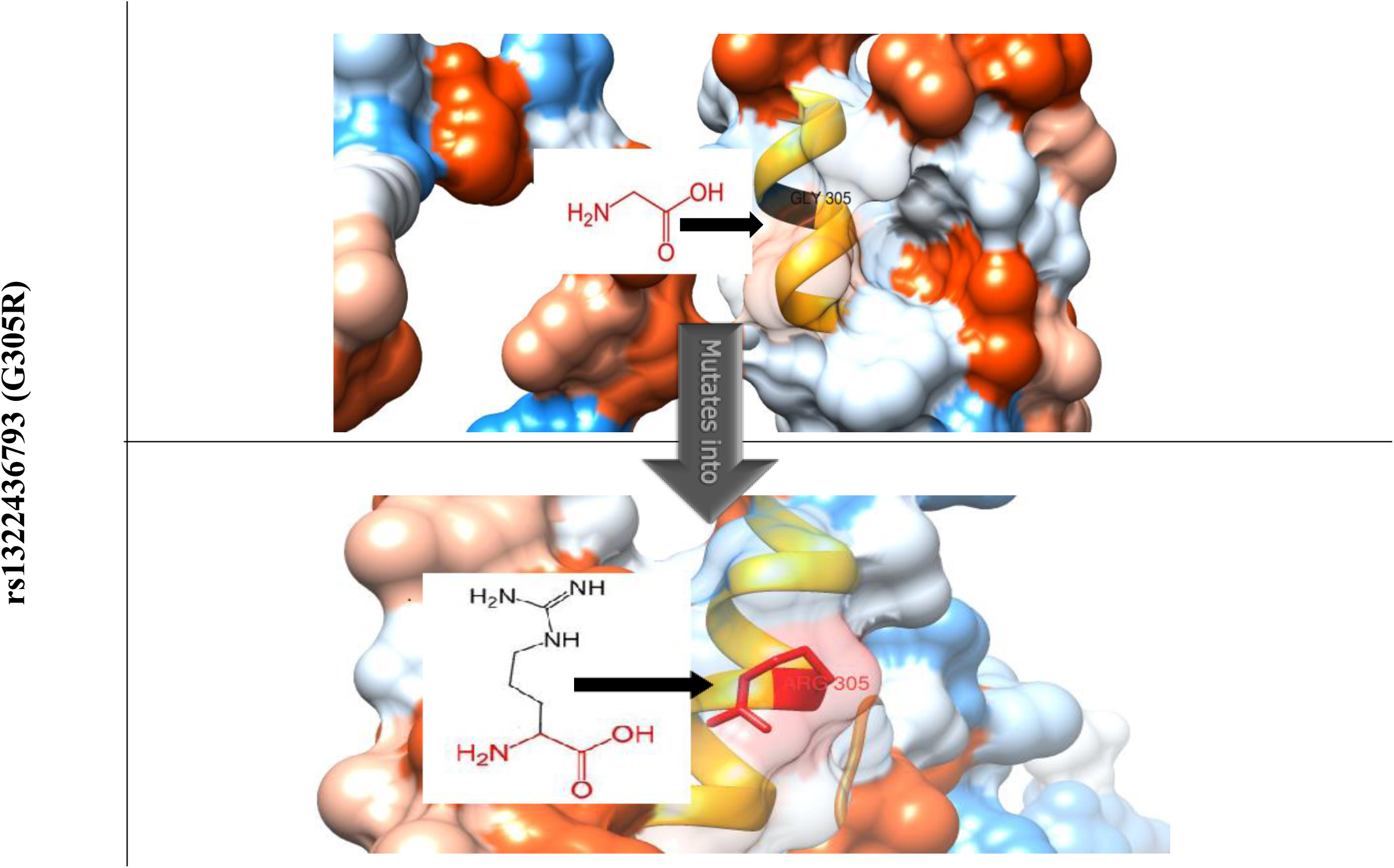
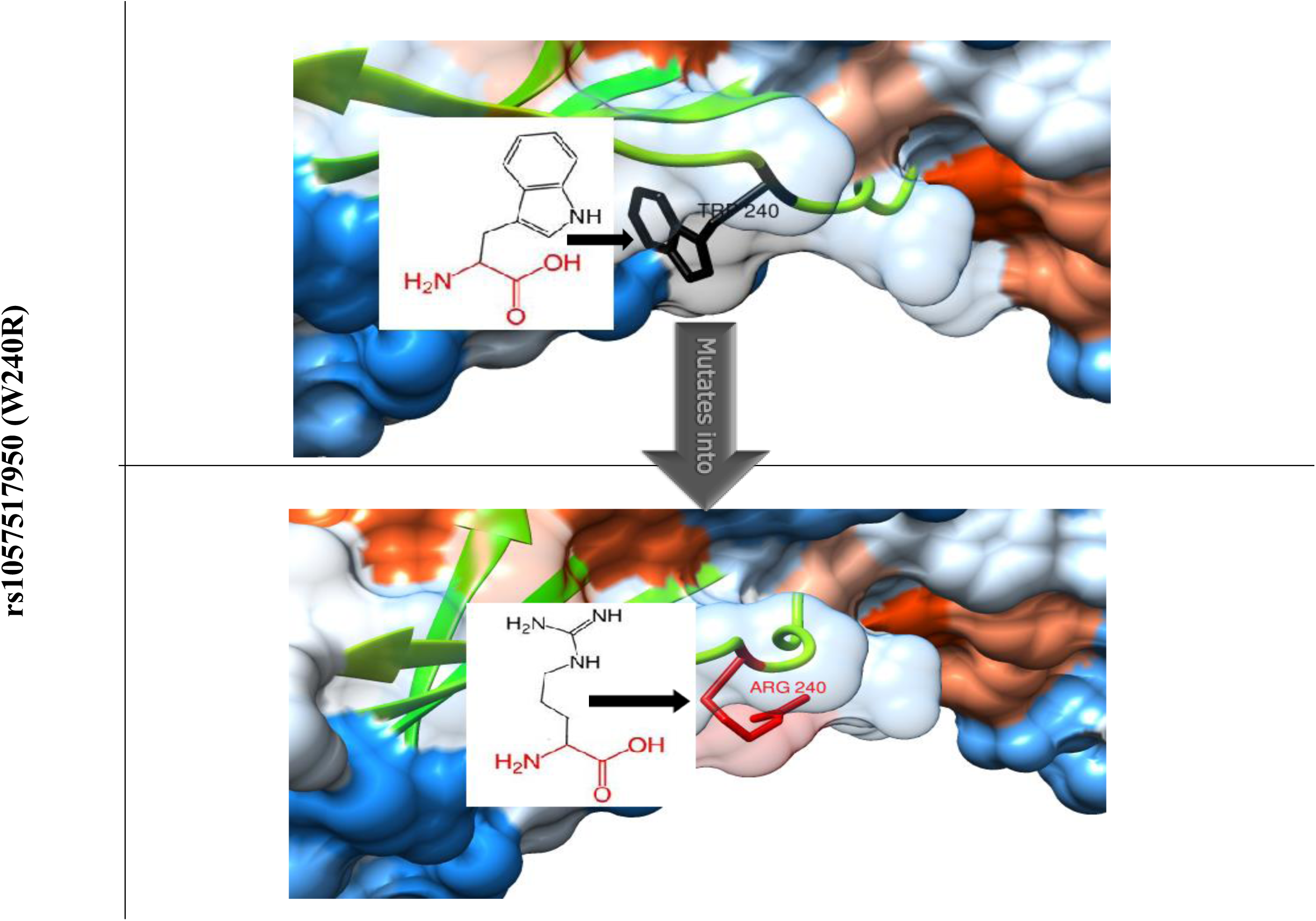
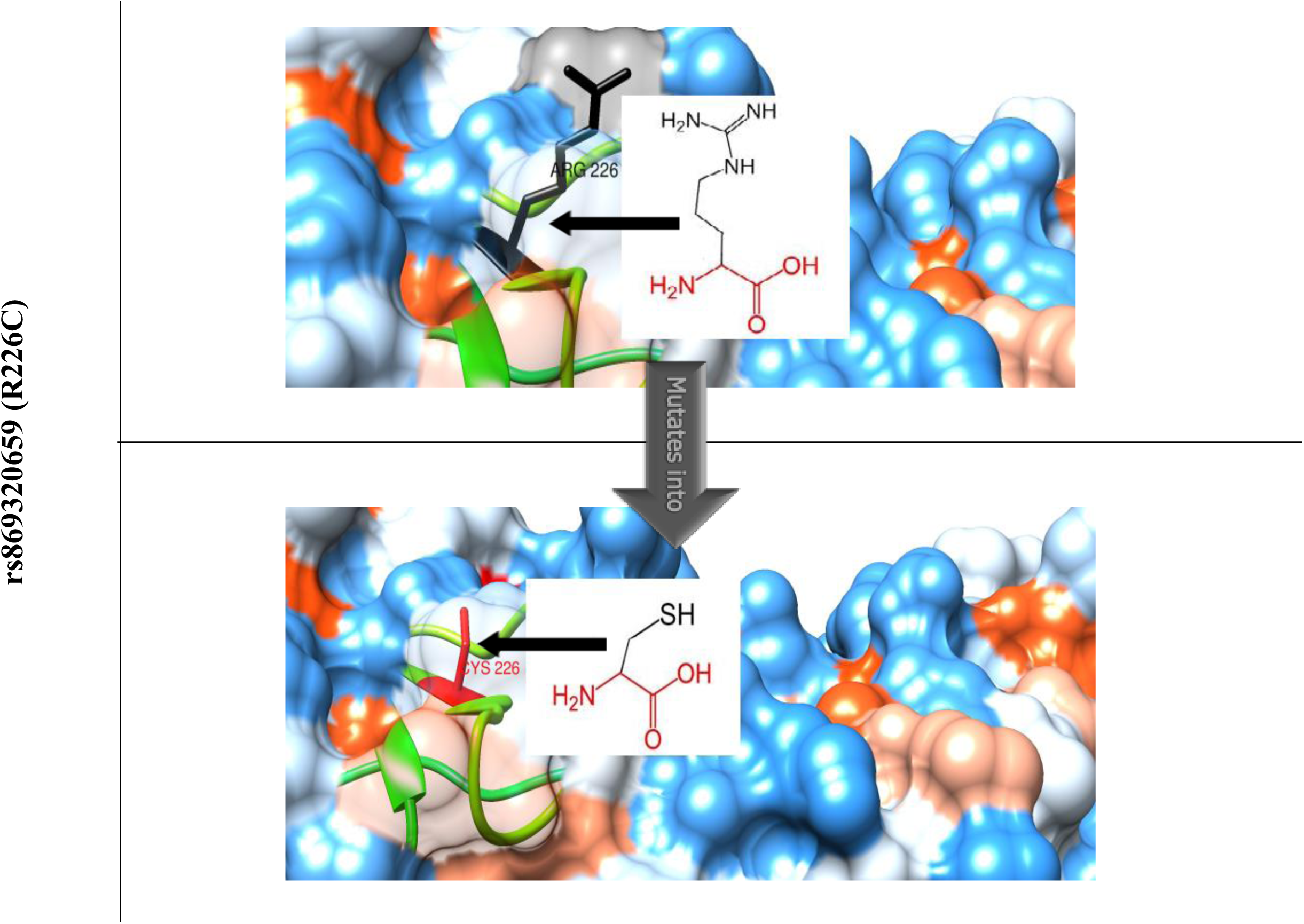
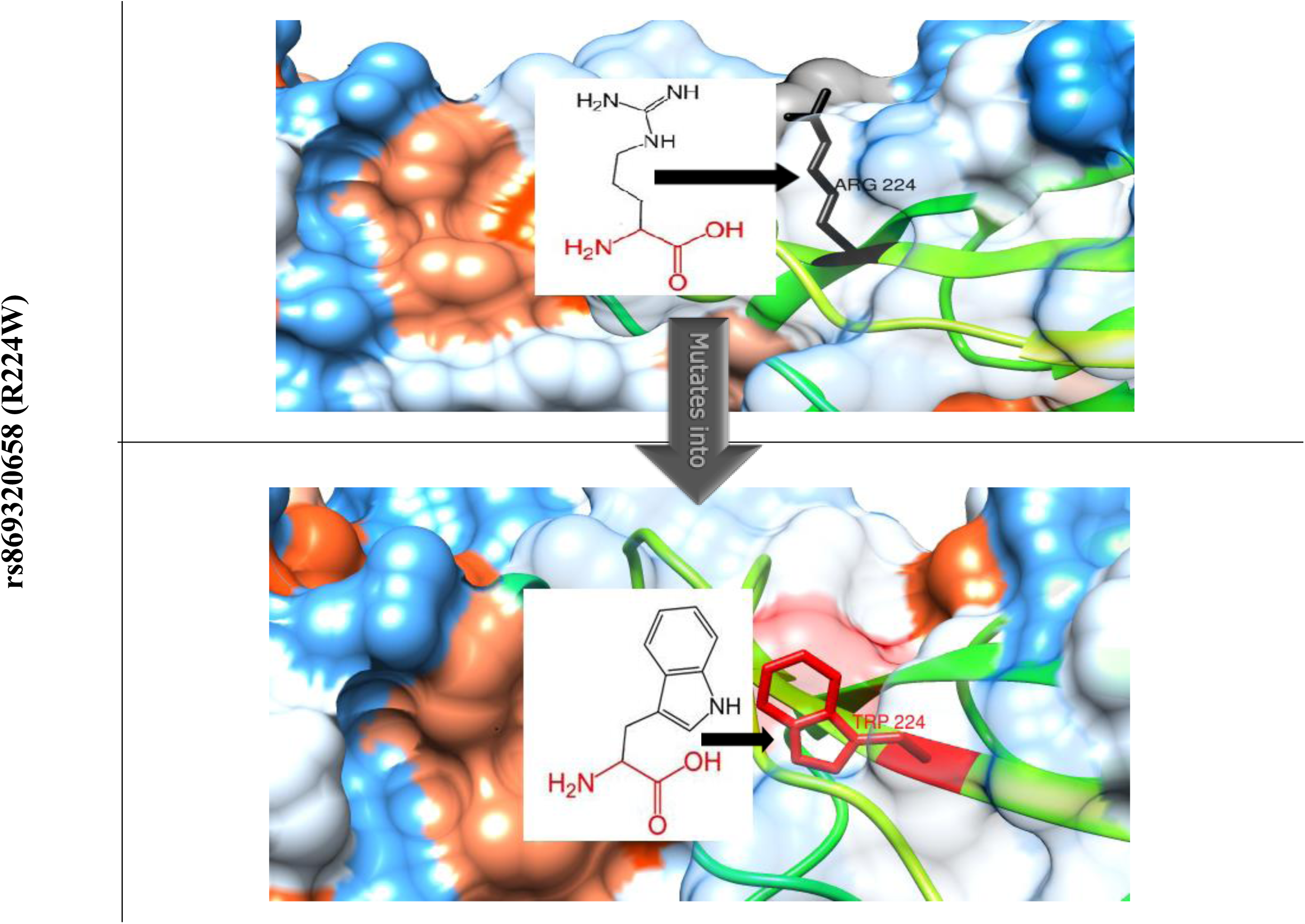
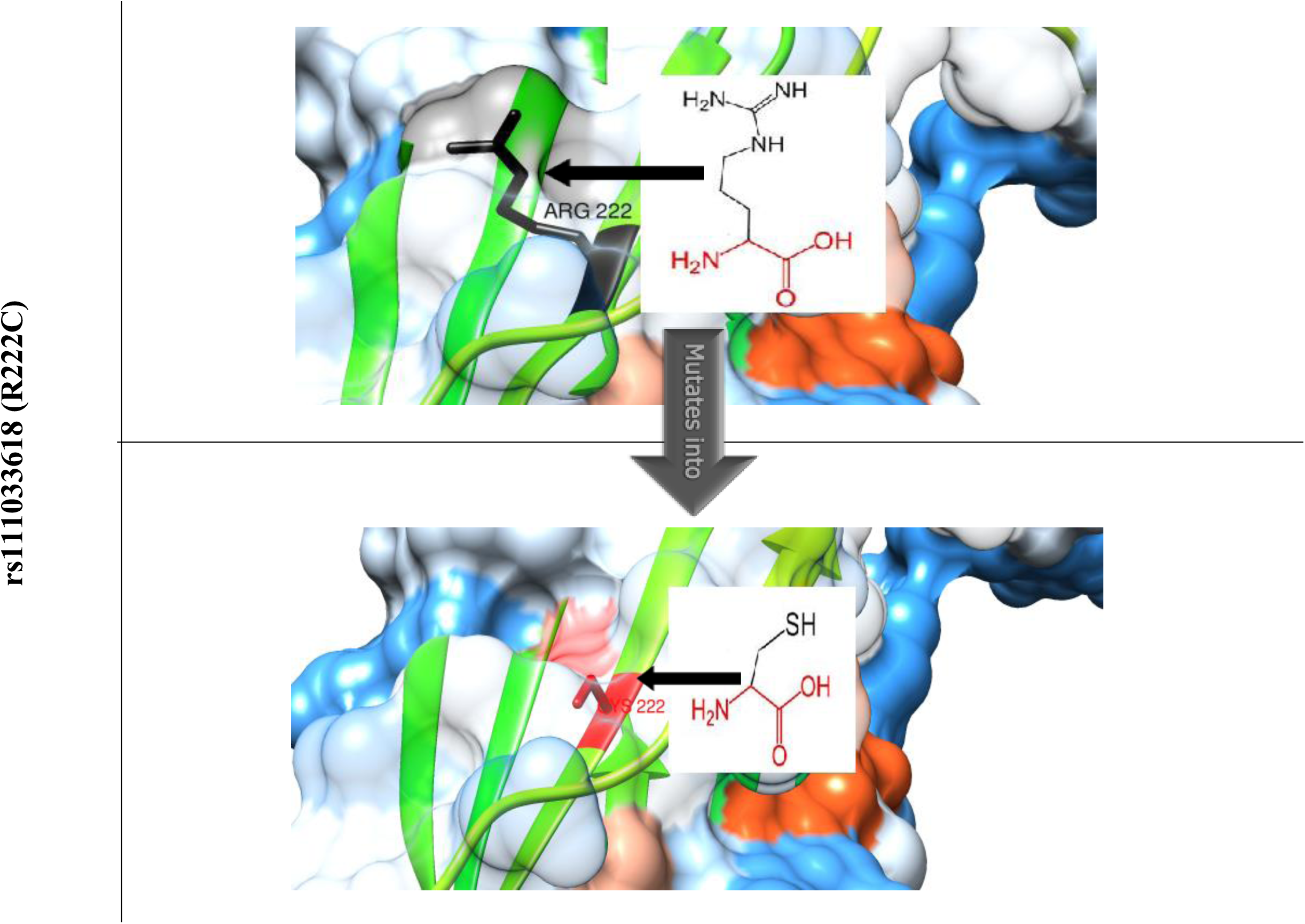
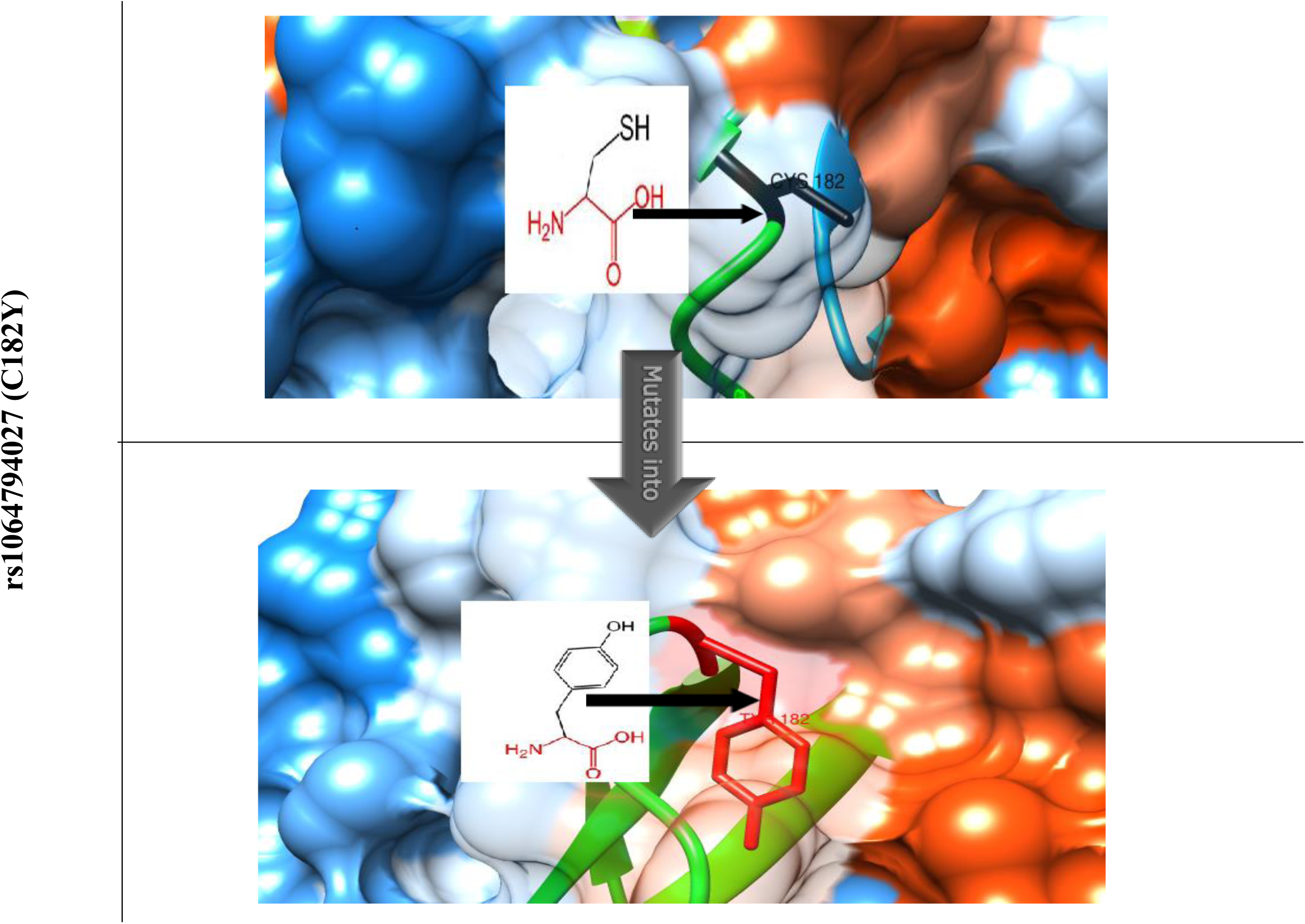
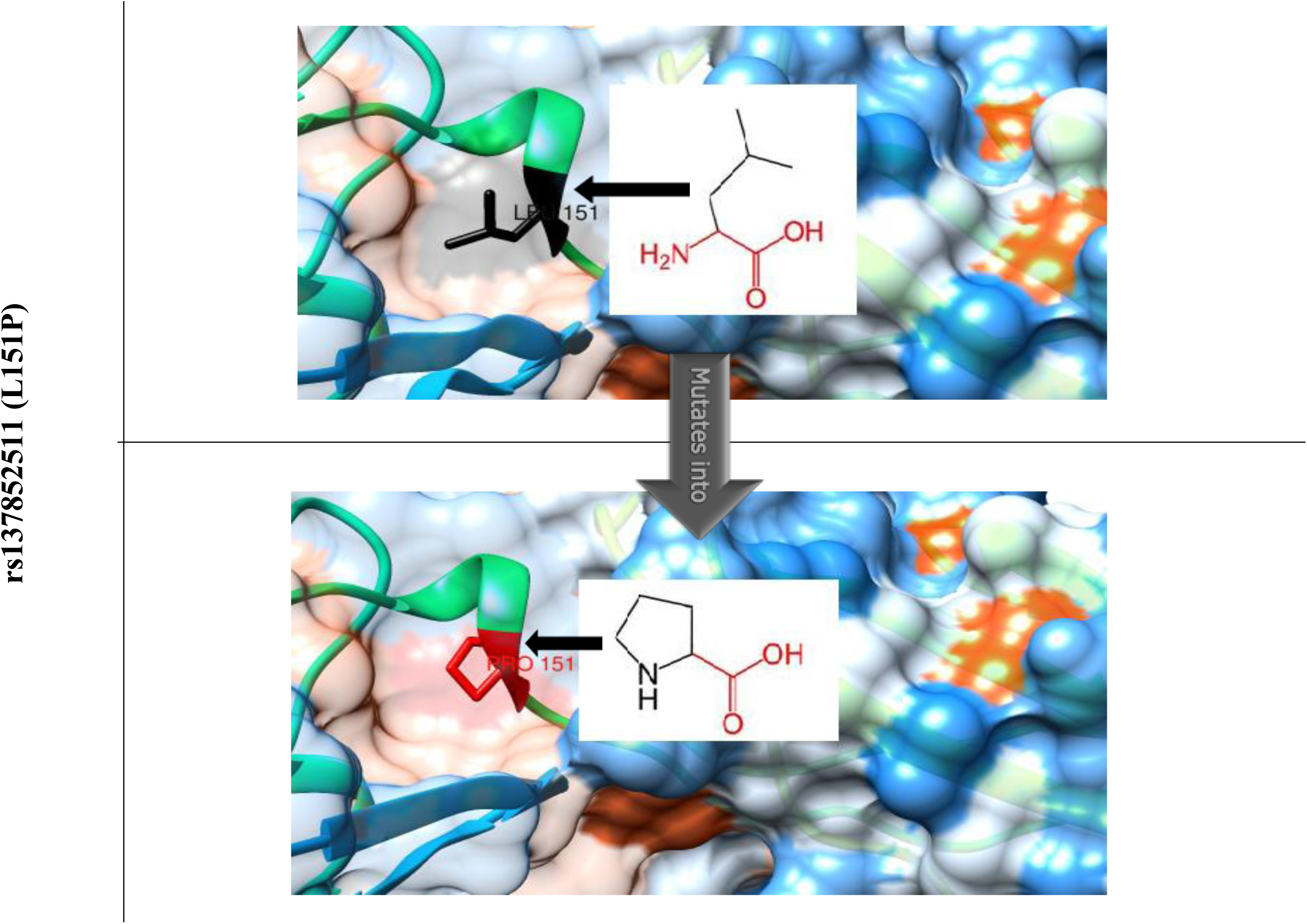
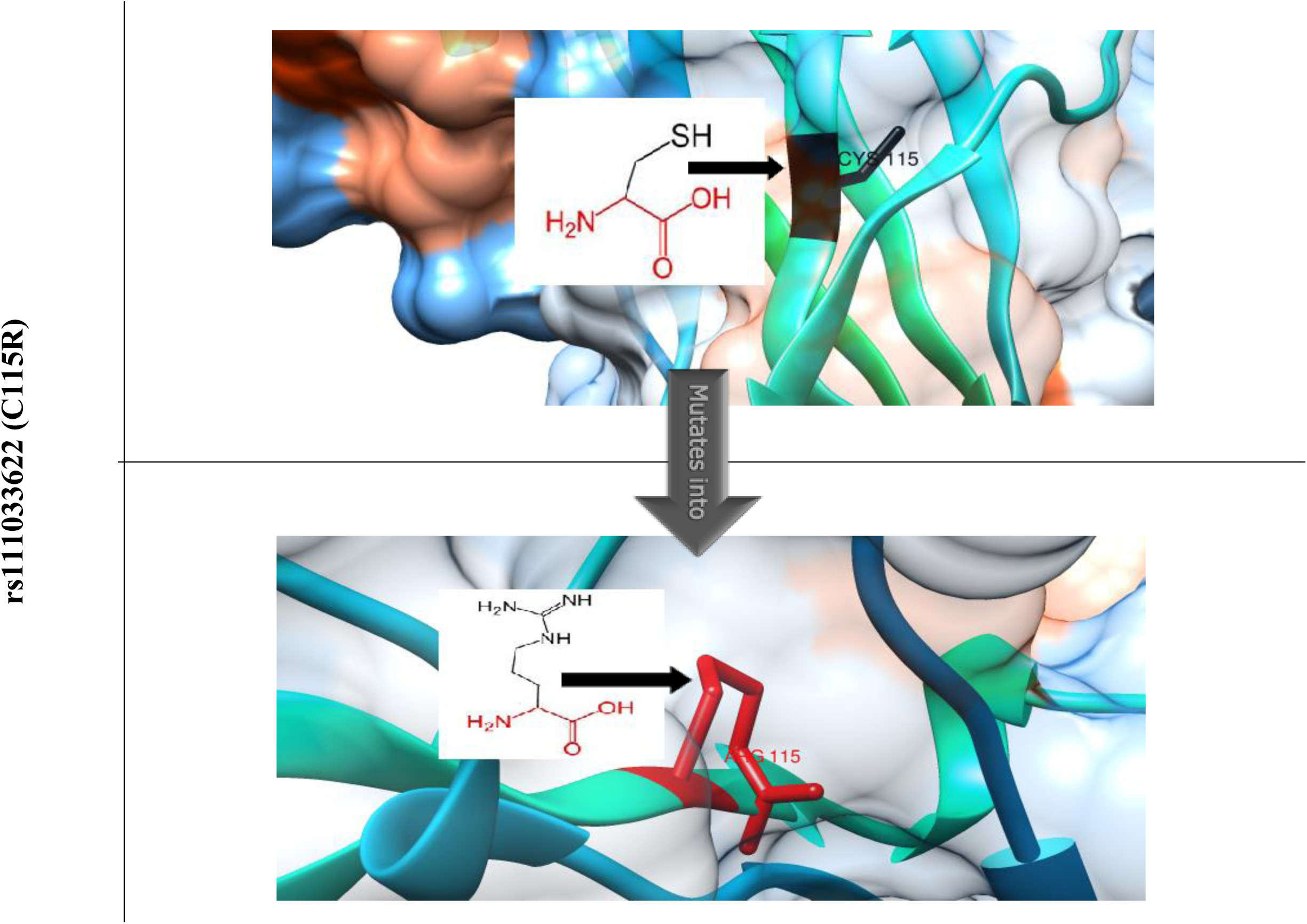
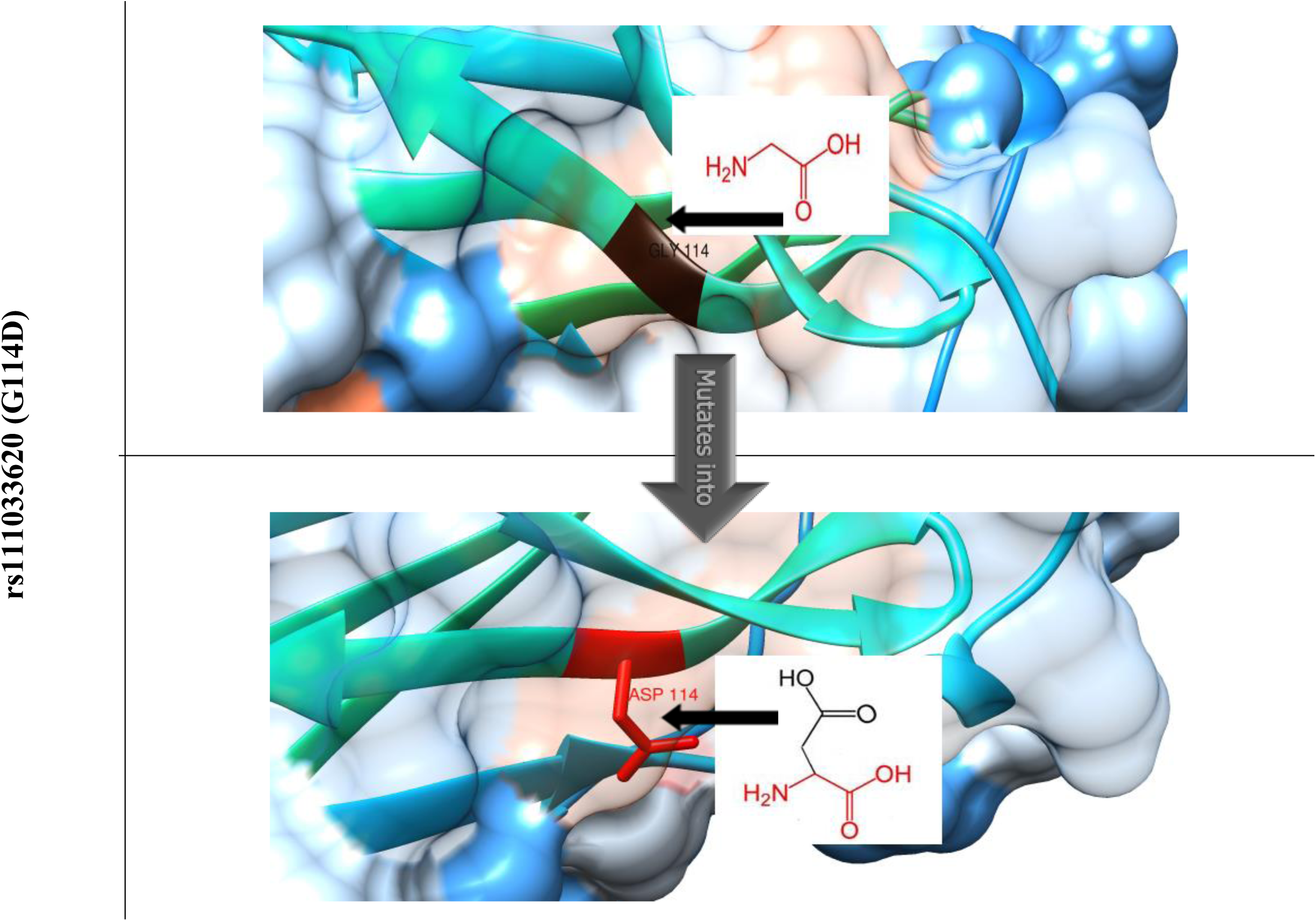
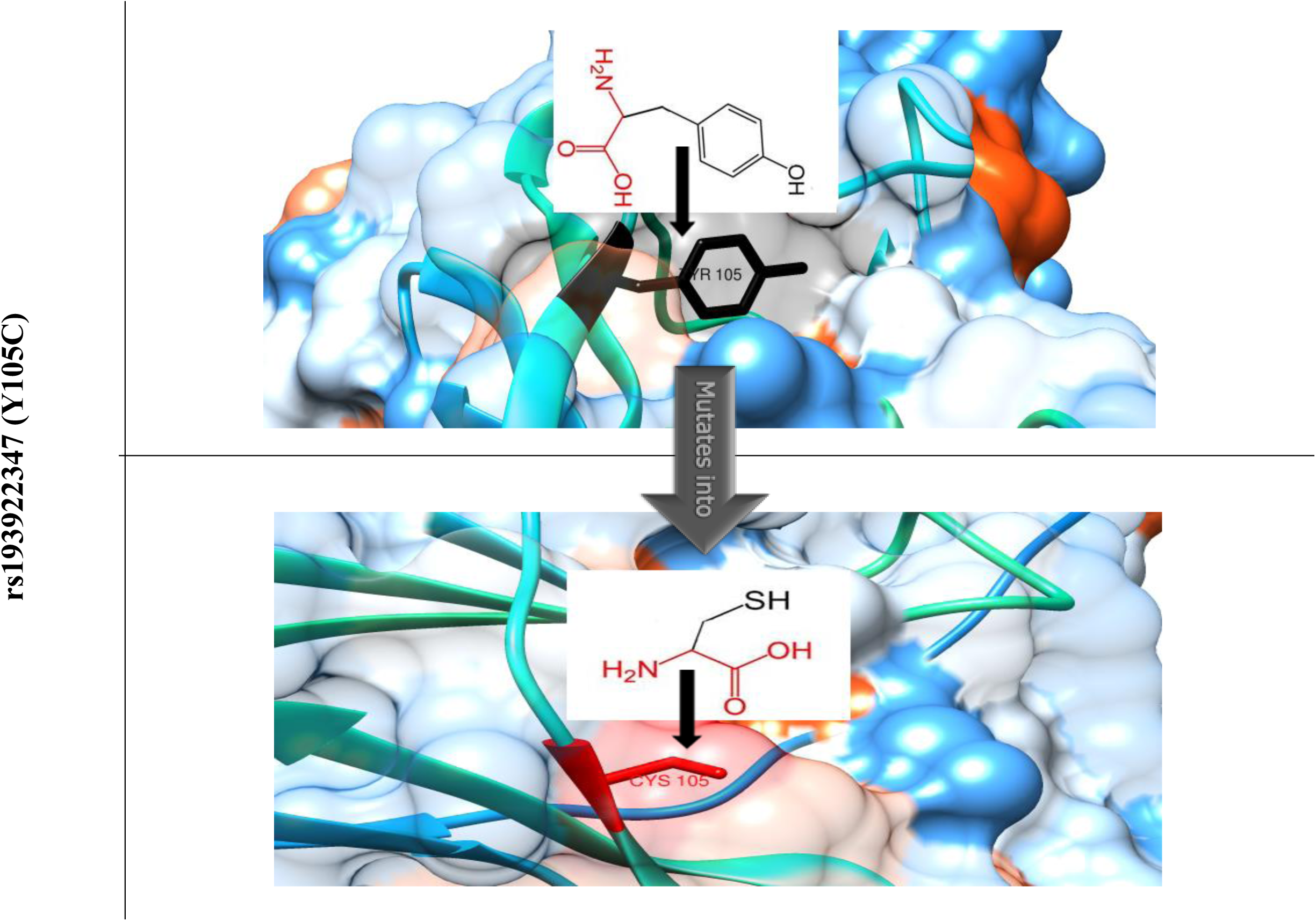
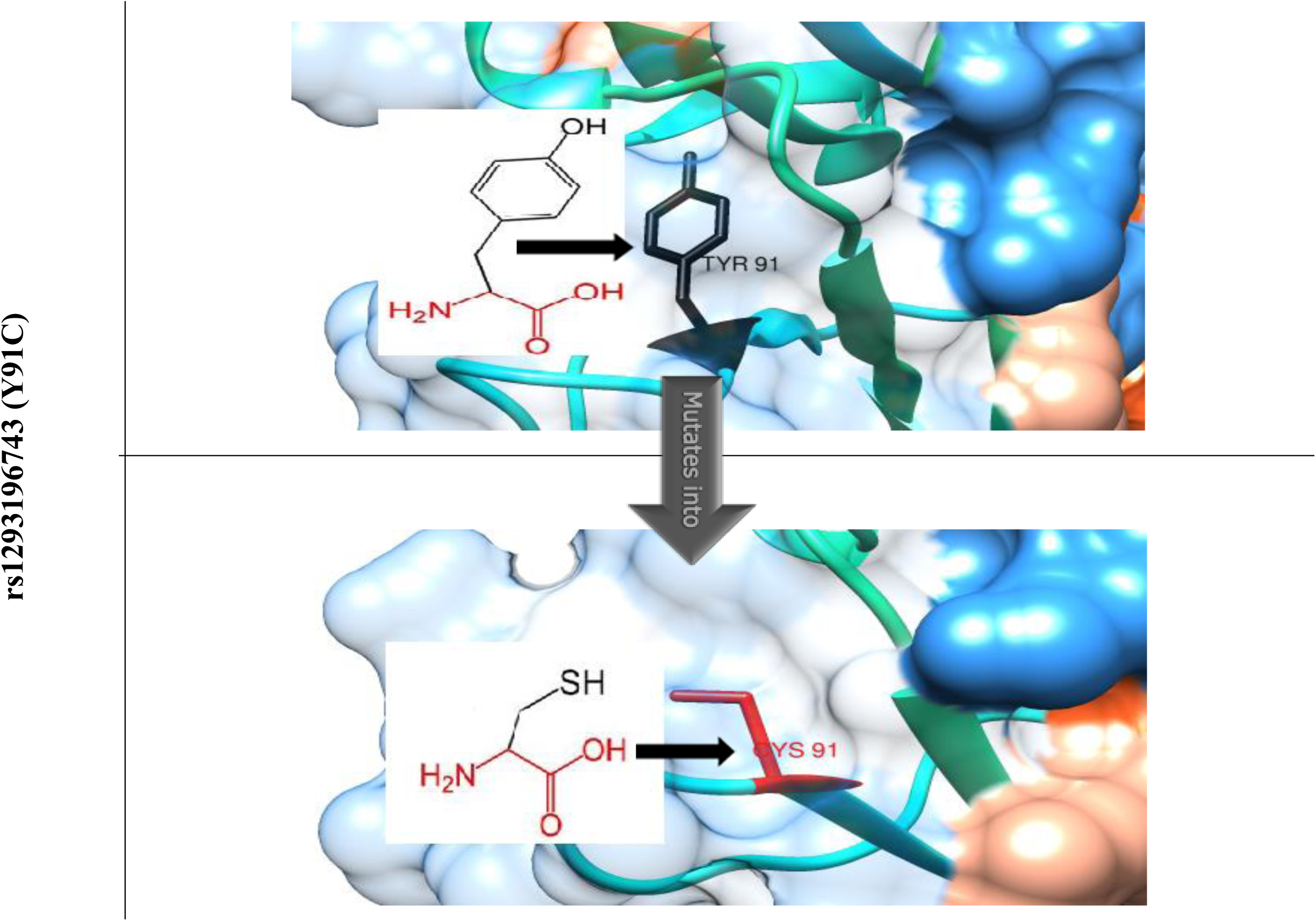
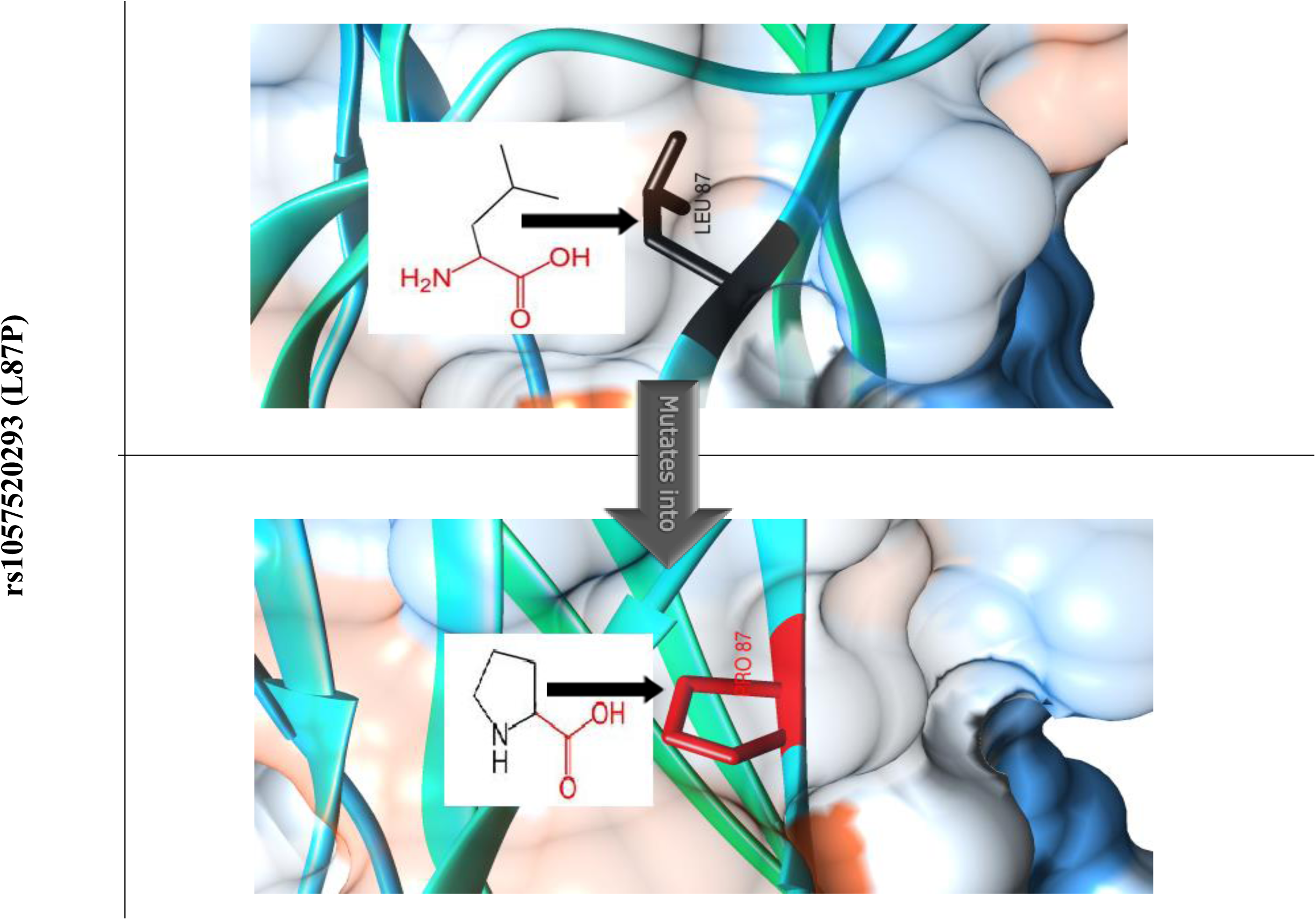
Modeling of wild and Mutant Structure by Project HOPE and Chimera:

### 3.6.

**Table (4).**
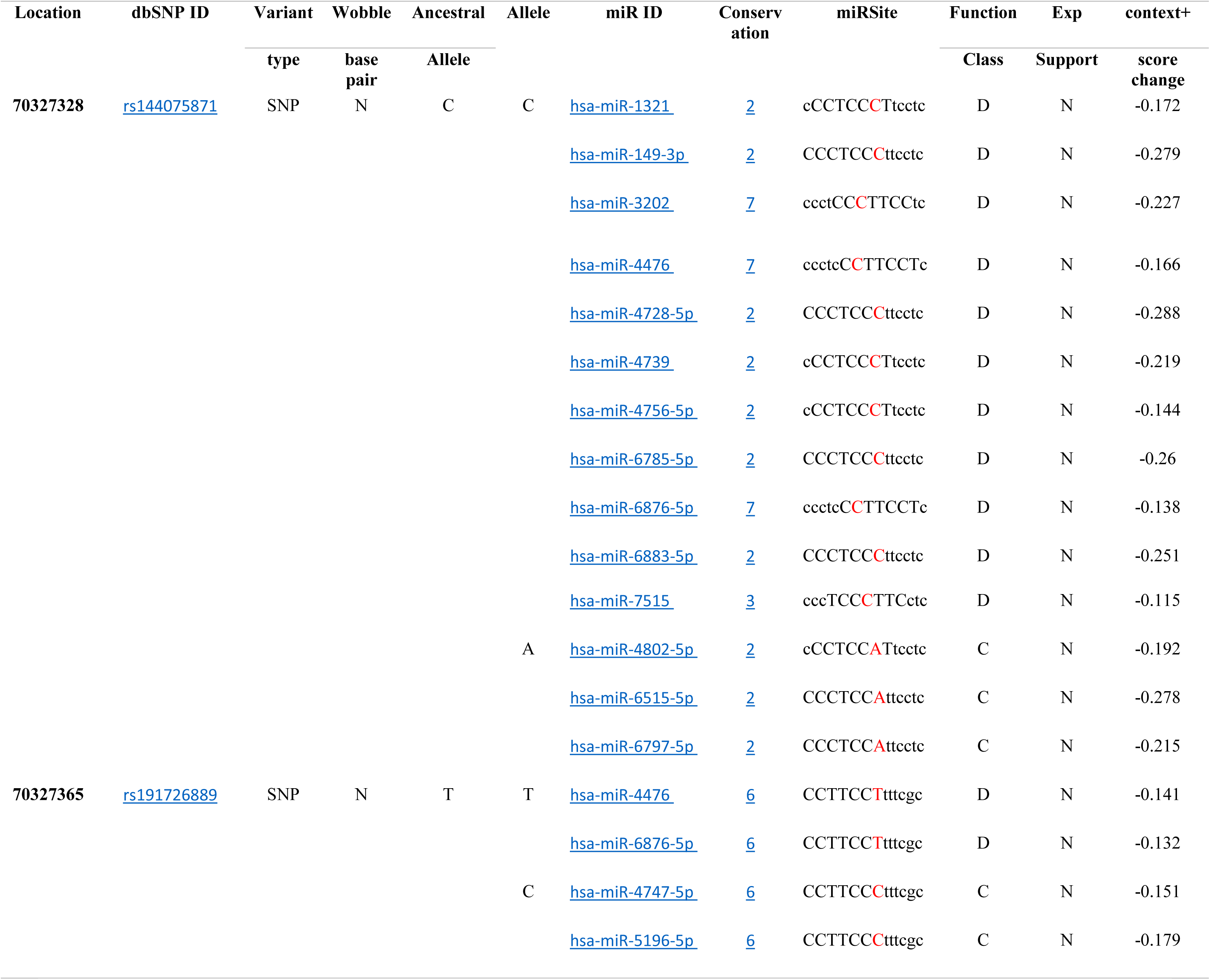
prediction of SNPs at the 3’UTR Region using PolymiRTS:

### 3.7. IL2RG Gene Interactions and network predicted by Genemania

**Figure 4:**
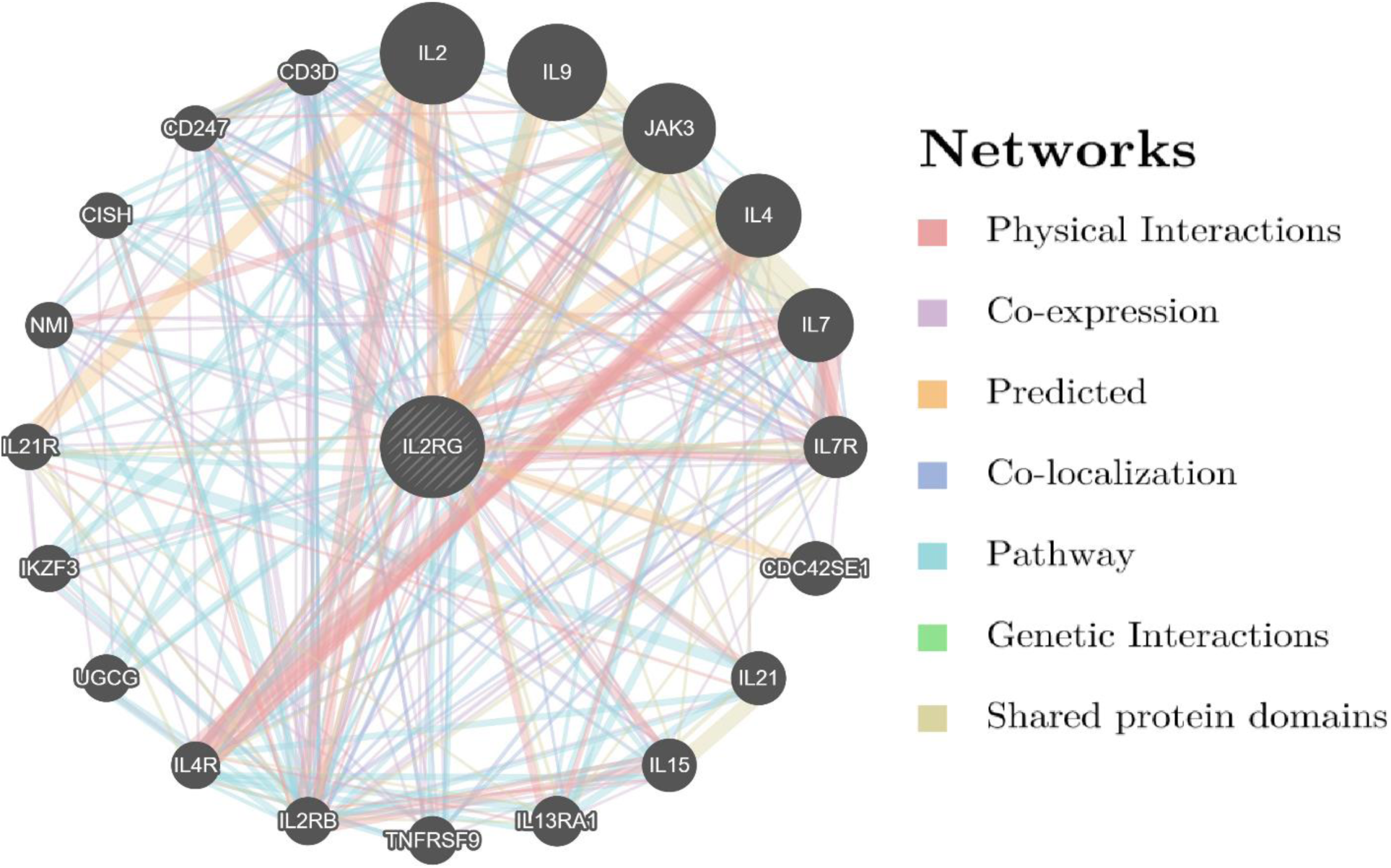
interactions between IL2RG and its related genes with paths colored according to their functions.

#### 3.7.1

**Table (7).**
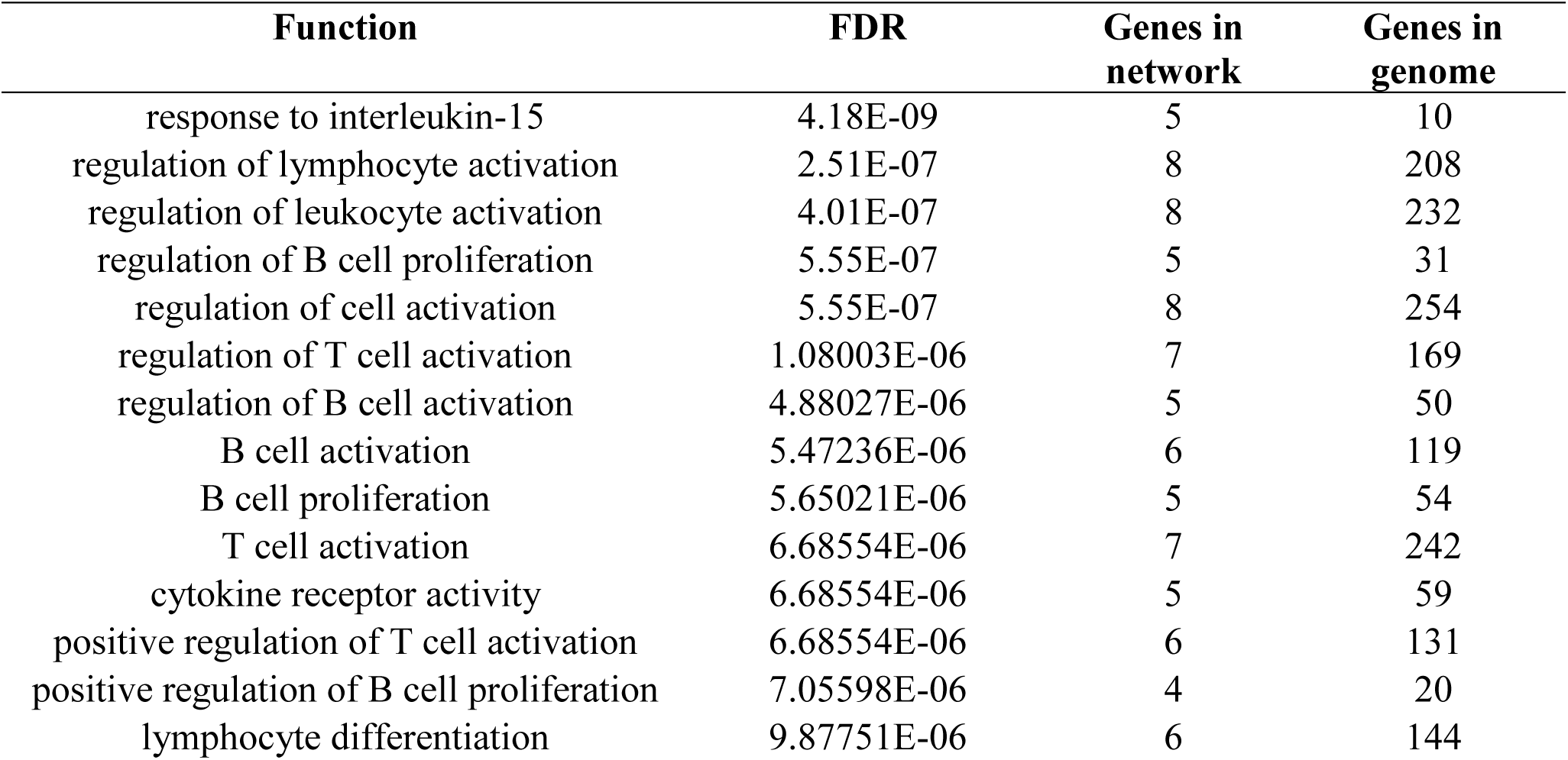
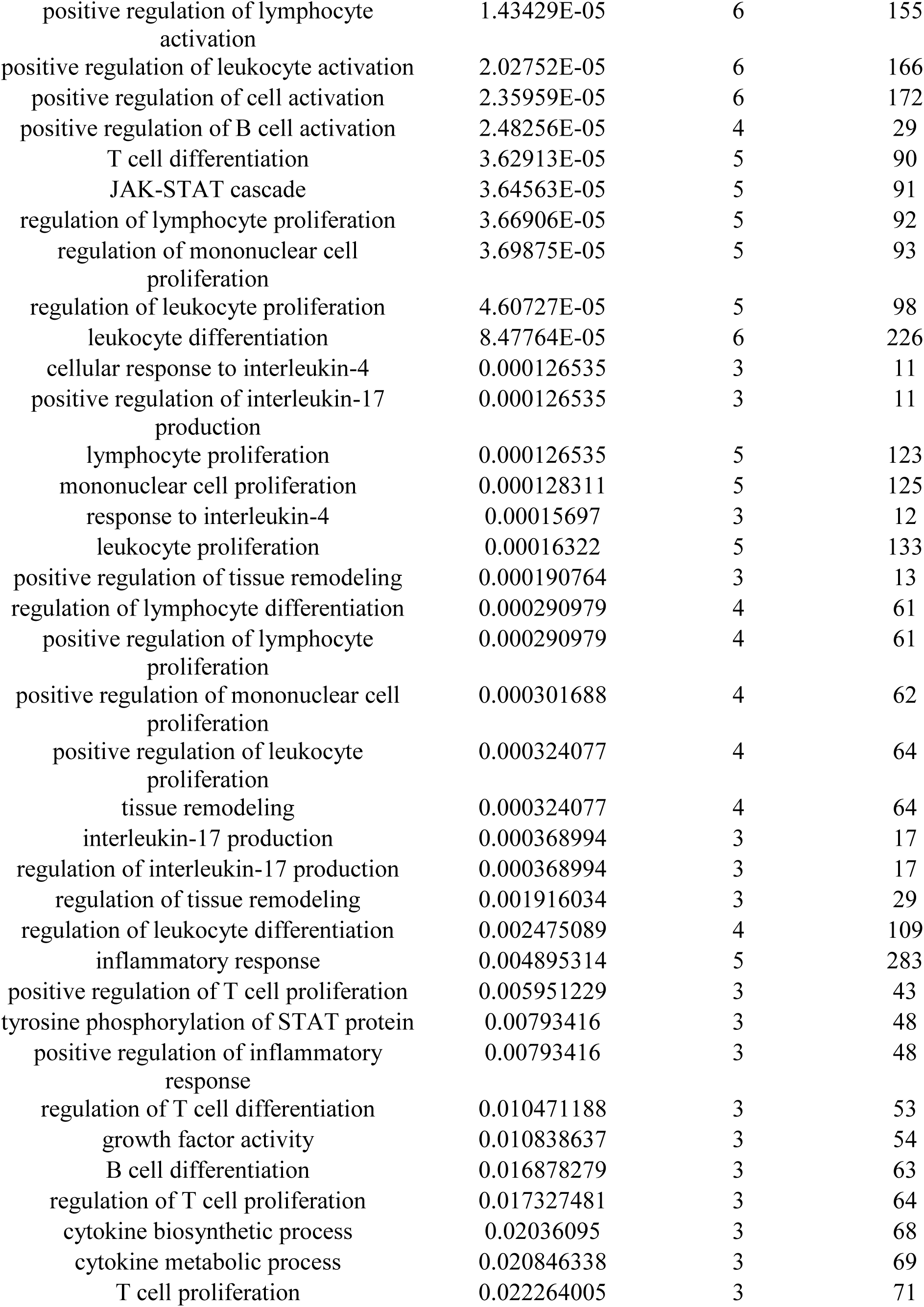
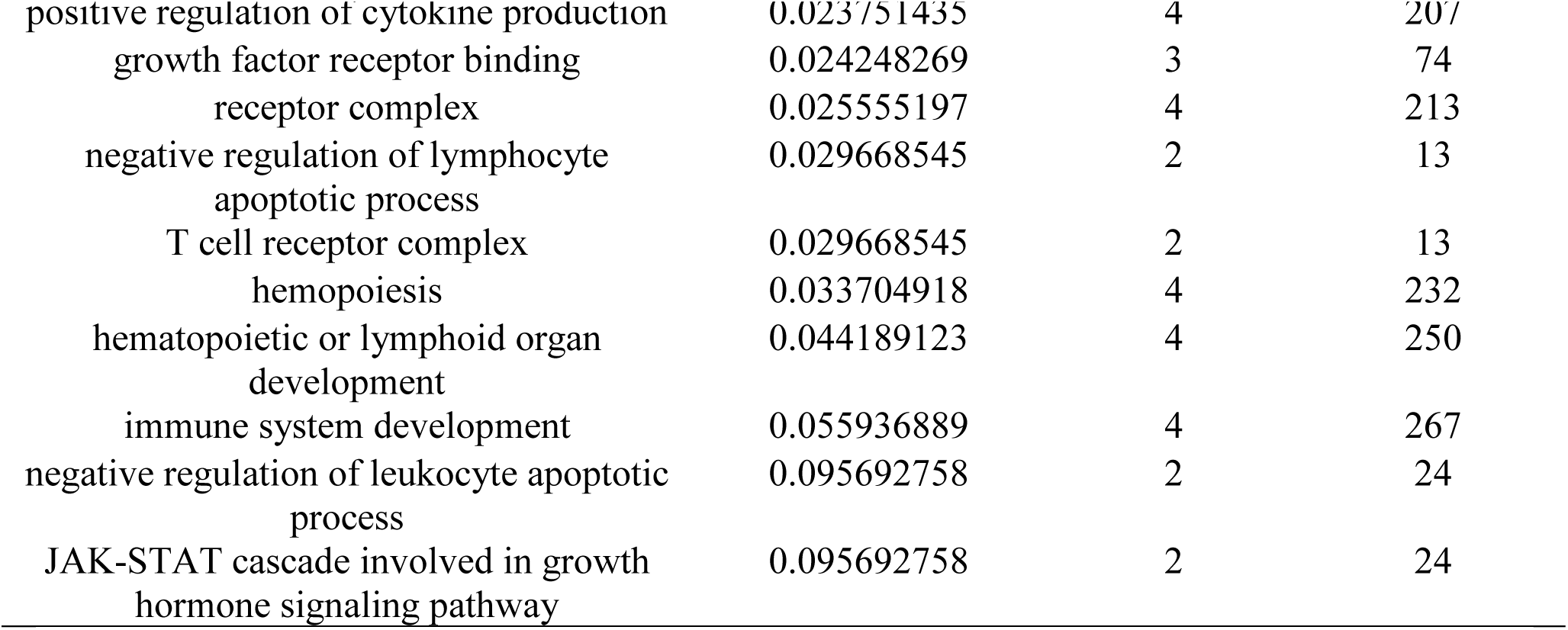
The IL2RG gene functions and its appearance in network and genome:

| Function | FDR | Genes in network | Genes in genome |
| --- | --- | --- | --- |
| response to interleukin-15 | 4.18E-09 | 5 | 10 |
| regulation of lymphocyte activation | 2.51E-07 | 8 | 208 |
| regulation of leukocyte activation | 4.01E-07 | 8 | 232 |
| regulation of B cell proliferation | 5.55E-07 | 5 | 31 |
| regulation of cell activation | 5.55E-07 | 8 | 254 |
| regulation of T cell activation | 1.08003E-06 | 7 | 169 |
| regulation of B cell activation | 4.88027E-06 | 5 | 50 |
| B cell activation | 5.47236E-06 | 6 | 119 |
| B cell proliferation | 5.65021E-06 | 5 | 54 |
| T cell activation | 6.68554E-06 | 7 | 242 |
| cytokine receptor activity | 6.68554E-06 | 5 | 59 |
| positive regulation of T cell activation | 6.68554E-06 | 6 | 131 |
| positive regulation of B cell proliferation | 7.05598E-06 | 4 | 20 |
| lymphocyte differentiation | 9.87751E-06 | 6 | 144 |
| positive regulation of lymphocyte activation | 1.43429E-05 | 6 | 155 |
| positive regulation of leukocyte activation | 2.02752E-05 | 6 | 166 |
| positive regulation of cell activation | 2.35959E-05 | 6 | 172 |
| positive regulation of B cell activation | 2.48256E-05 | 4 | 29 |
| T cell differentiation | 3.62913E-05 | 5 | 90 |
| JAK-STAT cascade | 3.64563E-05 | 5 | 91 |
| regulation of lymphocyte proliferation | 3.66906E-05 | 5 | 92 |
| regulation of mononuclear cell proliferation | 3.69875E-05 | 5 | 93 |
| regulation of leukocyte proliferation | 4.60727E-05 | 5 | 98 |
| leukocyte differentiation | 8.47764E-05 | 6 | 226 |
| cellular response to interleukin-4 | 0.000126535 | 3 | 11 |
| positive regulation of interleukin-17 production | 0.000126535 | 3 | 11 |
| lymphocyte proliferation | 0.000126535 | 5 | 123 |
| mononuclear cell proliferation | 0.000128311 | 5 | 125 |
| response to interleukin-4 | 0.00015697 | 3 | 12 |
| leukocyte proliferation | 0.00016322 | 5 | 133 |
| positive regulation of tissue remodeling | 0.000190764 | 3 | 13 |
| regulation of lymphocyte differentiation | 0.000290979 | 4 | 61 |
| positive regulation of lymphocyte proliferation | 0.000290979 | 4 | 61 |
| positive regulation of mononuclear cell proliferation | 0.000301688 | 4 | 62 |
| positive regulation of leukocyte proliferation | 0.000324077 | 4 | 64 |
| tissue remodeling | 0.000324077 | 4 | 64 |
| interleukin-17 production | 0.000368994 | 3 | 17 |
| regulation of interleukin-17 production | 0.000368994 | 3 | 17 |
| regulation of tissue remodeling | 0.001916034 | 3 | 29 |
| regulation of leukocyte differentiation | 0.002475089 | 4 | 109 |
| inflammatory response | 0.004895314 | 5 | 283 |
| positive regulation of T cell proliferation | 0.005951229 | 3 | 43 |
| tyrosine phosphorylation of STAT protein | 0.00793416 | 3 | 48 |
| positive regulation of inflammatory response | 0.00793416 | 3 | 48 |
| regulation of T cell differentiation | 0.010471188 | 3 | 53 |
| growth factor activity | 0.010838637 | 3 | 54 |
| B cell differentiation | 0.016878279 | 3 | 63 |
| regulation of T cell proliferation | 0.017327481 | 3 | 64 |
| cytokine biosynthetic process | 0.02036095 | 3 | 68 |
| cytokine metabolic process | 0.020846338 | 3 | 69 |
| T cell proliferation | 0.022264005 | 3 | 71 |
| positive regulation of cytokine production | 0.023751435 | 4 | 207 |
| growth factor receptor binding | 0.024248269 | 3 | 74 |
| receptor complex | 0.025555197 | 4 | 213 |
| negative regulation of lymphocyte apoptotic process | 0.029668545 | 2 | 13 |
| T cell receptor complex | 0.029668545 | 2 | 13 |
| hemopoiesis | 0.033704918 | 4 | 232 |
| hematopoietic or lymphoid organ development | 0.044189123 | 4 | 250 |
| immune system development | 0.055936889 | 4 | 267 |
| negative regulation of leukocyte apoptotic process | 0.095692758 | 2 | 24 |
| JAK-STAT cascade involved in growth hormone signaling pathway | 0.095692758 | 2 | 24 |

#### 3.7.2

**Table (8).** The genes co-expressed, co localized, Physical Interacted and share a domain with IL2RG:

**3.7.2 Table (8).** The genes co-expressed, co localized, Physical Interacted and share a domain with IL2RG:
| Gene 1 | Gene 2 | Weight | Network group |
| --- | --- | --- | --- |
| NMI | IL2RG | 0.007142981 | Co-expression |
| CD247 | IL2RG | 0.011854455 | Co-expression |
| CD3D | IL2RG | 0.010006283 | Co-expression |
| IL7R | IL2RG | 0.015151113 | Co-expression |
| TNFRSF9 | IL2RG | 0.01611196 | Co-expression |
| CD247 | IL2RG | 0.012152404 | Co-expression |
| CD3D | IL2RG | 0.012304108 | Co-expression |
| TNFRSF9 | IL2RG | 0.014275057 | Co-expression |
| IL21R | IL2RG | 0.011859321 | Co-expression |
| CD247 | IL2RG | 0.01117192 | Co-expression |
| CD3D | IL2RG | 0.014610344 | Co-expression |
| IL7R | IL2RG | 0.003189589 | Co-expression |
| TNFRSF9 | IL2RG | 0.006582764 | Co-expression |
| IL2RB | IL2RG | 0.007769996 | Co-expression |
| IL4R | IL2RG | 0.00616813 | Co-expression |
| CD247 | IL2RG | 0.003073129 | Co-expression |
| CD3D | IL2RG | 0.002789903 | Co-expression |
| IL4R | IL2RG | 0.012734761 | Co-expression |
| IL7 | IL2RG | 0.007766109 | Co-expression |
| IL2RB | IL2RG | 0.006711766 | Co-expression |
| IKZF3 | IL2RG | 0.007105181 | Co-expression |
| IL21R | IL2RG | 0.003766455 | Co-expression |
| CD247 | IL2RG | 0.005185788 | Co-expression |
| CD3D | IL2RG | 0.00601814 | Co-expression |
| JAK3 | IL2RG | 0.008937181 | Co-expression |
| CDC42SE1 | IL2RG | 0.011040032 | Co-expression |
| IKZF3 | IL2RG | 0.009303174 | Co-expression |
| CD3D | IL2RG | 0.007054924 | Co-expression |
| CD247 | IL2RG | 0.008991863 | Co-expression |
| IL15 | IL2RG | 0.008256315 | Co-expression |
| IL4R | IL2RG | 0.007494276 | Co-expression |
| IL7 | IL2RG | 0.004436255 | Co-expression |
| IL7R | IL2RG | 0.004471675 | Co-expression |
| CD247 | IL2RG | 0.006826574 | Co-expression |
| CD3D | IL2RG | 0.003450207 | Co-expression |
| NMI | IL2RG | 0.010466014 | Co-expression |
| CD3D | IL2RG | 0.012631002 | Co-expression |
| IL7R | IL2RG | 0.010826766 | Co-expression |
| JAK3 | IL2RG | 0.011127411 | Co-expression |
| IL4R | IL2RG | 0.004267869 | Co-expression |
| IL15 | IL2RG | 0.009034835 | Co-localization |
| TNFRSF9 | IL2RG | 0.008663382 | Co-localization |
| CD247 | IL2RG | 0.010449201 | Co-localization |
| CD3D | IL2RG | 0.005962262 | Co-localization |
| IL2 | IL2RG | 0.007554005 | Pathway |
| IL9 | IL2RG | 0.1314403 | Pathway |
| JAK3 | IL2RG | 0.007377151 | Pathway |
| IL4 | IL2RG | 0.018344974 | Pathway |
| IL7 | IL2RG | 0.024201902 | Pathway |
| IL7R | IL2RG | 0.027118763 | Pathway |
| IL21 | IL2RG | 0.024172688 | Pathway |
| IL15 | IL2RG | 0.024201902 | Pathway |
| IL2RB | IL2RG | 0.008450603 | Pathway |
| IL4R | IL2RG | 0.015245179 | Pathway |
| UGCG | IL2RG | 0.044504892 | Pathway |
| IKZF3 | IL2RG | 0.03478117 | Pathway |
| IL21R | IL2RG | 0.03414414 | Pathway |
| NMI | IL2RG | 0.048489608 | Pathway |
| CISH | IL2RG | 0.029071633 | Pathway |
| CD247 | IL2RG | 0.008765063 | Pathway |
| CD3D | IL2RG | 0.01024757 | Pathway |
| IL2 | IL2RG | 0.008473313 | Pathway |
| JAK3 | IL2RG | 0.012780515 | Pathway |
| IL4 | IL2RG | 0.014036303 | Pathway |
| TNFRSF9 | IL2RG | 0.09801694 | Pathway |
| IL2RB | IL2RG | 0.012710958 | Pathway |
| IL4R | IL2RG | 0.02225277 | Pathway |
| UGCG | IL2RG | 0.041583646 | Pathway |
| IKZF3 | IL2RG | 0.035283282 | Pathway |
| CISH | IL2RG | 0.06464156 | Pathway |
| CD247 | IL2RG | 0.013865737 | Pathway |
| CD3D | IL2RG | 0.013662125 | Pathway |
| IL2 | IL2RG | 0.32064575 | Physical Interactions |
| JAK3 | IL2RG | 0.5392597 | Physical Interactions |
| IL7 | IL2RG | 0.2520761 | Physical Interactions |
| IL15 | IL2RG | 0.19908392 | Physical Interactions |
| IL2RB | IL2RG | 0.13526194 | Physical Interactions |
| IL2 | IL2RG | 0.48999068 | Physical Interactions |
| JAK3 | IL2RG | 0.047650374 | Physical Interactions |
| IL21 | IL2RG | 0.35071465 | Physical Interactions |
| IL13RA1 | IL2RG | 0.32248512 | Physical Interactions |
| IL4R | IL2RG | 0.0740218 | Physical Interactions |
| IL4 | IL2RG | 0.39940926 | Physical Interactions |
| IL7 | IL2RG | 0.4357728 | Physical Interactions |
| IL7R | IL2RG | 0.33060825 | Physical Interactions |
| IL7 | IL2RG | 0.25101426 | Physical Interactions |
| IL7R | IL2RG | 0.34089583 | Physical Interactions |
| IL2 | IL2RG | 0.066997014 | Physical Interactions |
| JAK3 | IL2RG | 0.03302985 | Physical Interactions |
| IL4 | IL2RG | 0.09594477 | Physical Interactions |
| IL7 | IL2RG | 0.22980283 | Physical Interactions |
| IL7R | IL2RG | 0.0603281 | Physical Interactions |
| IL15 | IL2RG | 0.16295333 | Physical Interactions |
| IL2RB | IL2RG | 0.03907459 | Physical Interactions |
| IL4R | IL2RG | 0.041646395 | Physical Interactions |
| IL9 | IL2RG | 0.5554309 | Predicted |
| CDC42SE1 | IL2RG | 0.37048033 | Predicted |
| IL2 | IL2RG | 0.6641241 | Predicted |
| IL4 | IL2RG | 0.6641241 | Predicted |
| JAK3 | IL2RG | 1 | Predicted |
| IL7R | IL2RG | 0.040768523 | Shared protein domains |
| IL13RA1 | IL2RG | 0.034895726 | Shared protein domains |
| IL2RB | IL2RG | 0.040768523 | Shared protein domains |
| IL4R | IL2RG | 0.04114318 | Shared protein domains |
| IL21R | IL2RG | 0.04076852 | Shared protein domains |
| IL7R | IL2RG | 0.016019475 | Shared protein domains |
| IL13RA1 | IL2RG | 0.06732253 | Shared protein domains |

## 4. Discussion

*IL2RG* gene was investigated in dbSNP National Center of Biotechnology Information (NCBI public database). This gene containing a total of 253 SNPs in coding region, of which 153 were missense, 85 synonymous, eight nonsense, seven frame shift and 71 were in the non-coding region, of which 50 in 3′un-translated region (3’ UTR) and 21 in 5’ un-translated region (5’UTR). We selected the missense coding SNPs and 3′UTR SNPs for our investigation.

Our study revealed that 12 nsSNPs in the coding region were predicted to be damaging by various software. Five of them were novel; rs1322436793(G305R), rs1064794027(C182Y), rs111033620(G114D), rs193922347(Y105C) and rs1293196743(Y91C). The remaining seven nsSNPs were described by previous study to be damaging and disease related; rs1057517950 (W240R), rs869320659 (R226C), rs869320658 (R224W), rs111033618 (R222C), rs137852511 (L151P), rs111033622 (C115R) and rs1057520293 (L87P). Our study was in agreement with Weiping Tan et al who detected the rs1057517950 (W240R) mutation in a Chinese family with X-SCID. Their 4-month-old boy with X-SCID has a single point homozygous missense mutation within *IL2RG* exon 5: c. 718 T > C, p. W240R. The results of *IL2RG* sequencing showed that his mother and two sisters were heterozygous for the mutation and thus were asymptomatic carriers, while his father’s *IL2RG* gene was normal ^(48)^. In 2013 a study conducted by Alsina L et al described the vertical transmission of the somatic rs869320659 (R226C) *IL2RG* mosaicism from the mother to the child as the genetic mechanism underlying the patient with X-SCID ^(49)^. In 1997 Sharfe N et al revealed that the rs111033618 (R222C) mutation leads to an atypical phenotype presentation of SCID with a normal thymus gland, mitogen response and normal numbers of T and B cells. However, despite the normal number of T cells, the T cells demonstrate a reduced ability to bind IL-2 and thus incomplete participation in antigenic responses ^(50)^. Another study reported that in the rs111033618 (R222C) mutation signaling of γc cytokines for developmental processes was sufficient but the peripheral differentiation and function of T lymphocytes, B lymphocytes and NK cells was significantly reduced. Consequently, this resulted in more severe clinical phenotype than expected from T cell counts ^(18)^. Several studies investigated the frequency and variety of *IL2RG* mutations that cause SCID detected the mutations that had been predicted by our study to be damaging; rs1057517950 (W240R) ^(51)^, rs869320659 (R226C) ^(7, 51-53)^ and rs869320658 (R224W) ^(7, 51, 53)^. rs137852511 (L151P) mutation was detected in a 5-year-old boy by Speckmann C et al. This mutation was found in B, NK, and epithelial cells, whereas the gene sequence T cells were normal, indicating reversion of the mutation in a common T-cell precursor. This genetic correction in T cells resulted in restoration of immune function and an a typical phenotype with mild immunodeficiency ^(54)^. rs111033622 (C115R) was described in a one-year old boy with atypical X-SCID. The results of genetic analysis revealed normal expression of the γc chain and an absence of the γc gene mutation in the T cells. Reversion of the mutation in early T-cell precursors in this patient led to partially functional lymphocyte clones ^(24)^. rs1057520293 (L87P) mutation had been described in X-SCID patient with mild phenotype and delayed diagnosis due to the reversion in a common lymphoid progenitor ^(55)^.

Two SNPs out of 50 at the 3’UTR were predicted to disrupt miRNAs binding sites and hence affect the gene expression. rs144075871 SNP and rs191726889 SNP contained (D) functional classes that disrupted a conserved miRNA site and (C) functional class that created a new site of miRNA.

In rs1322436793 (G305R) mutation the mutant residue is bigger than the wild-type residue, this might lead to bumps. The wild-type residue charge is neutral while the mutant residue charge is positive, this can cause repulsion of ligands or other residues with the same charge. The wild-type residue is more hydrophobic than the mutant residue. Moreover in both rs1322436793 (G305R) and rs111033620 (G114D) mutations the torsion angles for this residue are unusual. Only glycine is flexible enough to make these torsion angles, mutation into another residue will force the local backbone into an incorrect conformation and will disturb the local structure. The wild-type residue in rs1064794027 (C182Y) and rs111033620 (G114D) mutations was buried in the core of the protein, the mutant residue is bigger probably will not fit. The wild-type residue is more hydrophobic than the mutant residue. The mutation will cause loss of hydrophobic interactions in the core of the protein. The mutant residue in rs193922347 (Y105C) and rs1293196743 (Y91C) mutations is smaller than the wild-type residue. This will cause a possible loss of external interactions in (Y105C) and will cause an empty space in the core of the protein in (Y91C). The mutant residue is more hydrophobic than the wild-type residue. The mutated residue is located in a domain that is important for binding of other molecules and in contact with residues in a domain that is also important for binding. The mutation might disturb the interaction between these two domains and as such affect the function of the protein.

All eleven mutations (G → R, W → R, R → C, R → W, R → C, C → Y, L → P,G → D, Y → C, Y → C) predicted a decrease of the protein stability, while only mutation (C → R) predicted an increase of stability of protein. *IL2RG* gene function and activities illustrated by GENEMAN which showed that *IL2RG* gene has many functions such as cellular response to interleukin-4, cytokine receptor activity, response to interleukin-15, response to interleukin-4. *IL2RG* gene was predicted to be related to 20 genes (*CDC42SE1, IL9, TNFRSF9, IL21, IL4, IL13RA1, IL15, IKZF3, IL2, NMI, JAK3, UGCG, IL7, CISH, IL21R, ILR7, CD247, CD3D, IL2RB,* and *IL4R*).

This study is computational and has limitations; in vivo genetic analysis is recommended. This will improve our understanding of how complex human phenotype is inherited. SNPs revealed by this study could be considered as important candidates in causing diseases related to *IL2RG* mutation and could be used as diagnostic markers. We hope our results will provide useful information that needed to help researchers to do further studies and also may help in diagnosis and genetic screening of SCID

## 5. Conclusions

Computational analysis of SNPs has become a very valuable tool in order to discriminate neutral SNPs from SNPs of likely functional importance and associated with disease. In this study we used different software to predict the damaging mutations of *IL2RG* gene; 12 nsSNPs were predicted by different software to be the most damaging mutations for *IL2RG* protein altering its size, charge, hydrophobicity, stability and physiochemical properties and thus leading to loss of the protein’s function. Out of the 12 nsSNPs, 5 nsSNPs were novel and the rest were described by other studies to be disease related. Two SNPs out of 50 at the 3’UTR were predicted to disrupt miRNAs binding sites and hence affect the gene expression. This might result in alteration of the gene function.

## 6. Acknowledgment

The authors desire to acknowledge the excited cooperation of Africa City of Technology-Khartoum, Sudan.

